# Linking single-cell measurements of mass, growth rate, and gene expression

**DOI:** 10.1101/331686

**Authors:** Robert J. Kimmerling, Sanjay M. Prakadan, Alejandro J. Gupta, Nicholas L. Calistri, Mark M. Stevens, Selim Olcum, Nathan Cermak, Riley S. Drake, Alex K. Shalek, Scott R. Manalis

## Abstract

We introduce a microfluidic platform that enables single-cell mass and growth rate measurements upstream of single-cell RNA-sequencing (scRNA-seq) to generate paired single-cell biophysical and transcriptional data sets. Biophysical measurements are collected with a serial suspended microchannel resonator platform (sSMR) that utilizes automated fluidic state switching to load individual cells at fixed intervals, achieving a throughput of 120 cells per hour. Each single-cell is subsequently captured downstream for linked molecular analysis using an automated collection system. From linked measurements of a murine leukemia (L1210) and pro-B cell line (FL5.12), we identify gene expression signatures that correlate significantly with cell mass and growth rate. In particular, we find that both cell lines display a cell-cycle signature that correlates with cell mass, with early and late cell-cycle signatures significantly enriched amongst genes with negative and positive correlations with mass, respectively. FL5.12 cells also show a significant correlation between single-cell growth efficiency and a G1-S transition signature, providing additional transcriptional evidence for a phenomenon previously observed through biophysical measurements alone. Importantly, the throughput and speed of our platform allows for the characterization of phenotypes in dynamic cellular systems. As a proof-of-principle, we apply our system to characterize activated murine CD8+ T cells and uncover two unique features of CD8+ T cells as they become proliferative in response to activation: i) the level of coordination between cell cycle gene expression and cell mass increases, and ii) translation-related gene expression increases and shows a correlation with single-cell growth efficiency. Overall, our approach provides a new means of characterizing the transcriptional mechanisms of normal and dysfunctional cellular mass and growth rate regulation across a range of biological contexts.

## Background

Recent experimental advancements have dramatically improved the throughput and cost-efficiency of single-cell RNA-sequencing (scRNA-seq) [1–3]. However, gene expression measurements alone only provide a portion of the information necessary to characterize complex cellular processes [4, 5]. Thus, parallel efforts have focused on linking gene expression measurements with complementary single-cell data that can provide further information to help guide analyses and contextualize distinct cellular states. For instance, various multi-omic methods have been developed to link measurements such as protein abundance, or DNA sequence or methylation, with gene expression from the same single cell [6–9]. Gene expression measurements have also been linked to single-cell location within a tissue to enable the study of cellular development and differentiation at unprecedented detail [10–12]. Additionally, single-cell functional assays have been coupled with expression data to obtain novel insights into the relationships among cellular electrophysiology, morphology, and transcription [13]. Taken together, these approaches demonstrate the value of linked single-cell data sets to afford a deep understanding of various cellular functions and states that may be difficult to obtain through transcriptomic measurements alone.

Linked gene expression data sets are of particular interest when considering recent technological developments that have enabled the precise measurement of various single-cell biophysical properties, such as mass and growth rate [14, 15]. As highly integrative metrics of cellular state, these parameters offer unique insights into a wide range of biological phenomena, including: i) basic patterns of single-cell mass and growth regulation; ii) biophysical changes associated with immune cell activation; and, iii) heterogeneity of single-cell drug response in various cancers [16–18]. However, the approaches previously used to collect these biophysical measurements have precluded linking these properties with molecular information collected from the same cell.

Padovan-Merhar et al. recently reported key progress towards this goal, describing an imaging-based approach that allows for the enumeration of various transcripts that can be linked with single-cell volumetric measurements [19]. This work revealed novel insights into the relationship between transcript abundance and cell volume throughout the cell cycle. However, these imaging-based approaches are limited to measurements of cell size alone, have a low throughput, and have been coupled primarily with hybridization-based approaches that are limited in the total number of genes that can be measured in any given cell. To our knowledge, there have been no methods reported to date that allow for linked measurements of cellular mass, growth rate, and transcriptome-wide gene expression from the same cell. It has therefore been challenging to characterize the underlying transcriptional mechanisms responsible for the cellular mass and growth rate variability observed in a range of normal and dysfunctional biological contexts.

To address these limitations, we have developed a microfluidic platform that enables the measurement of single-cell mass and growth rate immediately upstream of scRNA-seq. Here, we describe this platform and demonstrate the reproducibility of the linked data sets it generates with a sufficient throughput to be applied to a broad range of dynamic biological contexts.

## Results and Discussion

### Serial SMR platform with downstream collection for scRNA-seq

Our system relies on a modified version of a serial suspended microchannel resonator (sSMR) device that has been described previously (Figure 1) [17]. Briefly, the sSMR utilizes an array of high-resolution singlecell buoyant mass sensors placed periodically along the length of a long microfluidic channel, allowing a single cell’s mass to be measured periodically as it traverses the channel. In addition to providing mass information, this series of measurements can also be used to determine the mass accumulation rate (MAR), or growth rate, of each cell. Real-time access to the data generated by each SMR mass sensor allows for peak detection in the final cantilever to be used as an indication of a cell exiting the mass sensor array; these peaks trigger the motion of a three-dimensional motorized stage to position a PCR tube containing lysis buffer to capture each single cell as it is flushed from the system (**Methods**).

**Figure 1.**
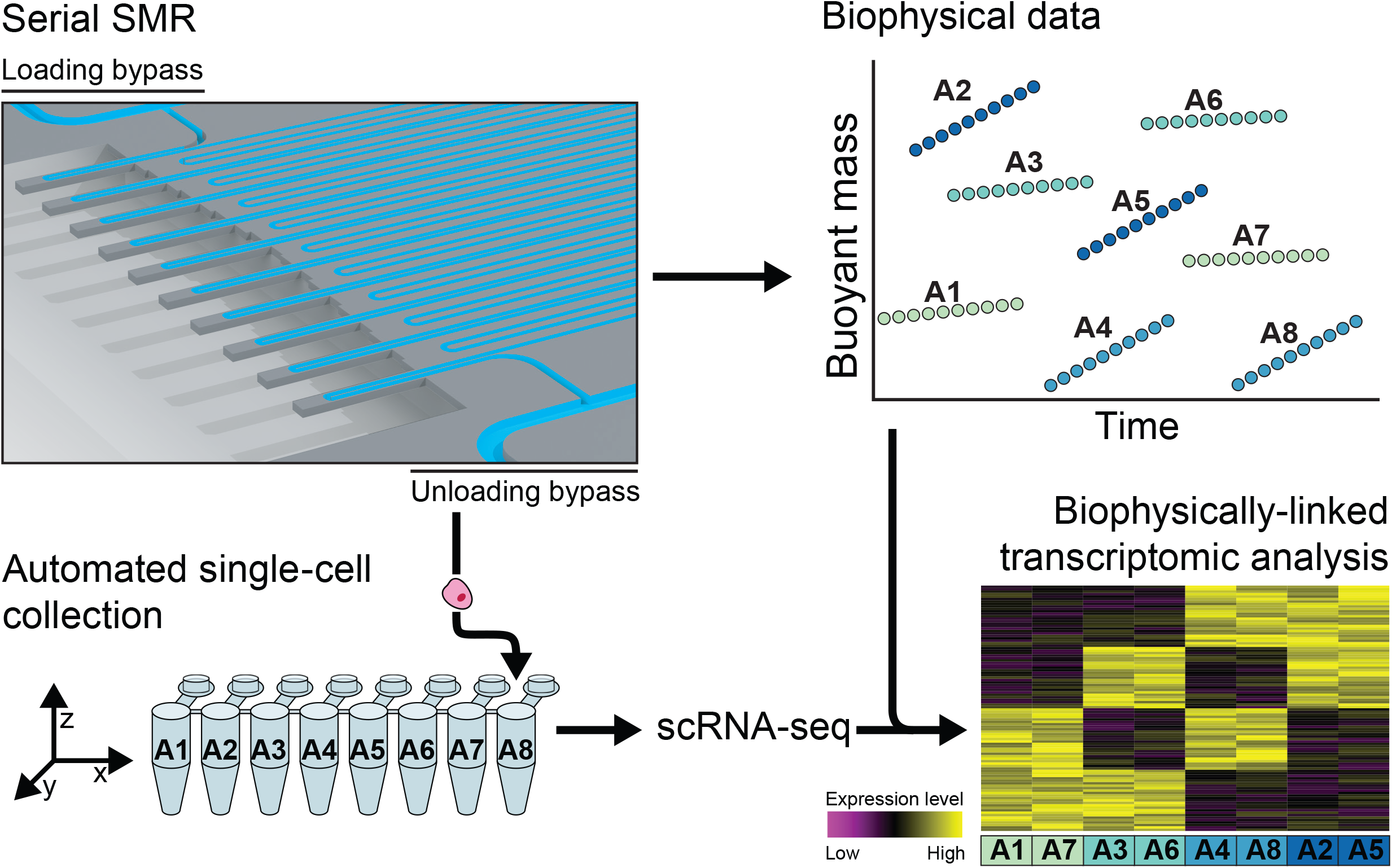
Serial SMR platform with downstream collection for scRNA-seq. Schematic representation of the serial SMR platform, which includes an array of SMR mass sensors, separated by a serpentine delay channel to periodically measure the buoyant mass of a single cell. Independent control of the upstream and downstream pressures applied to two bypass channels allows for single-cell spacing at the loading entrance of the array (top left of sSMR image) and single-cell isolation at the unloading exit (bottom right of sSMR image) (**Supplementary Figure 1, Supplementary Note 1**). Using real-time peak detection at the final mass sensor, a three-dimensional motorized stage is triggered to capture each individual cell directly in to lysis buffer for downstream single-cell RNA-sequencing. Based on well location, each cell is subsequently matched to its corresponding biophysical data collected from the sSMR including mass and MAR, as schematized in the top-right panel. These linked single-cell data sets can then be used to determine gene expression signatures associated with mass and growth rate variability, as schematized in the bottom-right panel.

The total time required to flush the system’s dead volume and release each single cell (20 seconds for the system implementation described here) sets a theoretical maximum throughput for the platform to avoid the collection of multiplets. Crucially, to minimize the frequency of failed capture events, we have implemented a new fluidic scheme whereby single cells are loaded into the array of mass sensors at fixed intervals (**Supplementary Figure 1, Supplementary Note 1**) [20]. Ultimately, this fluidic scheme allows us to achieve a throughput of one cell approximately every thirty seconds (for a throughput of up to 120 cells per hour) with minimal failed collection events due to co-release. This offers a two-fold throughput improvement over previous implementations of biophysical measurements alone, while offering the additional ability to capture each individual cell downstream for scRNA-seq.

### Unique gene expression profiles related to biophysical properties in two murine lymphoblast cell lines

To validate our method for collecting linked single-cell biophysical and gene expression data, we first measured two murine lymphoblast cell lines (L1210 and FL5.12) that have well-characterized mass and growth properties (Figure 2) [15–17]. Single cells collected downstream of the sSMR for scRNA-seq consistently yielded high-quality cDNA libraries, with 85 out of 87 individual L1210 cells and 124 out 144 individual FL5.12 cells with paired biophysical data passing initial quality controls (e.g., number of genes detected greater than 4,000, **Methods, Supplementary Figure 2**).

**Figure 2.**
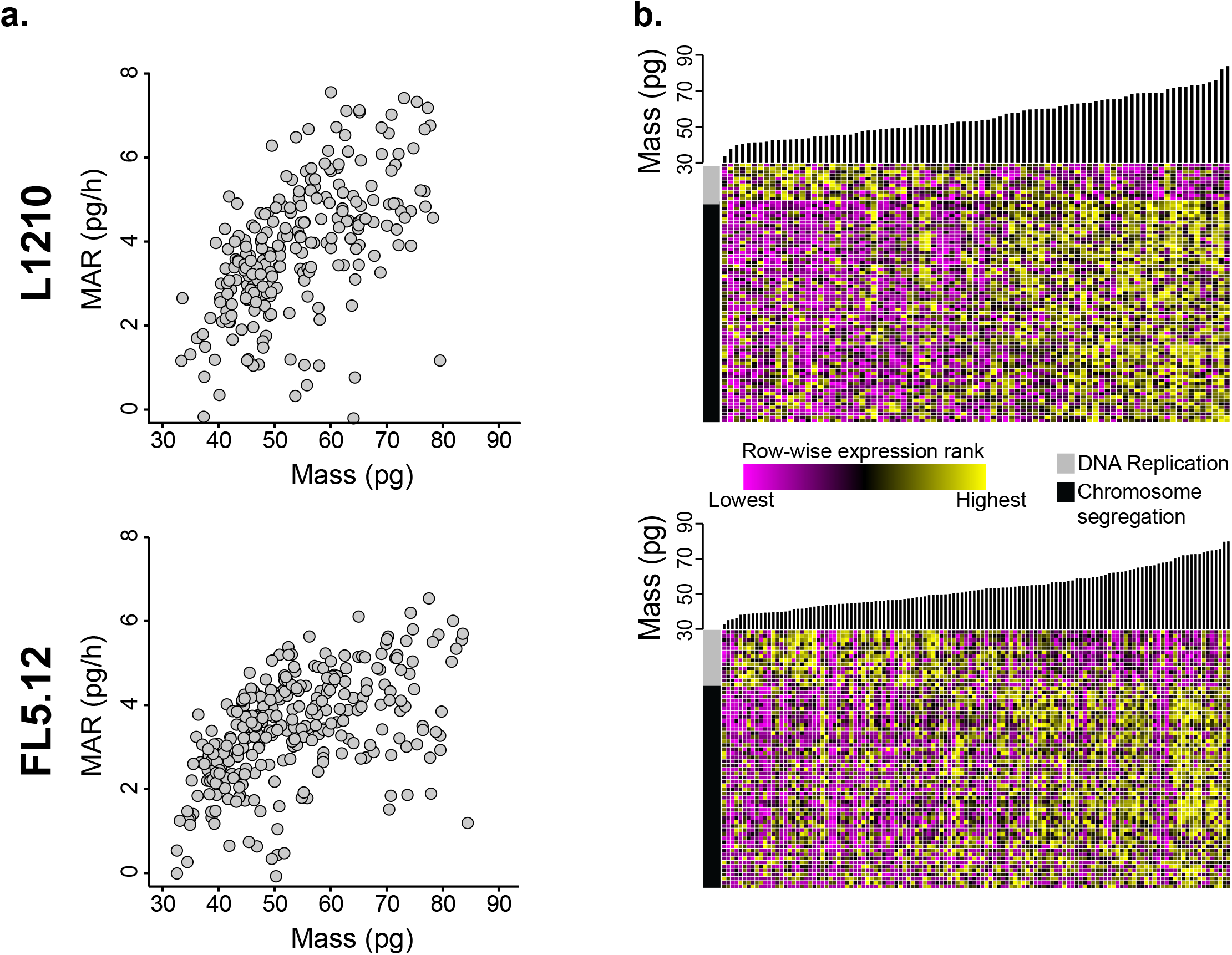
Linked biophysical and gene expression measurements of single L1210 and FL5.12 cells. (**a**) Plot of mass accumulation rate versus buoyant mass for single L1210 cells (top, n = 234) and single FL5.12 cells (bottom, n=296) measured in the sSMR. (**b**) Heat maps showing the relative expression of various cell cycle-related genes for subsets of the L1210 (top, n=85) and FL5.12 (bottom, n=124) cells depicted in (**a**) that were captured downstream for scRNA-seq. Cells are ordered by buoyant mass (bar plots above heat maps). Entries are colored by row-wise expression level rank where the cell with the highest expression level for a particular gene corresponds to yellow and the cell with the lowest expression level corresponds to magenta. As a demonstration, the heat map includes genes with expression levels that showed a significant correlation with buoyant mass from the chromosome segregation (black bar, n=66 and n=50 for the L1210 and FL5.12, respectively) and DNA replication (gray bar, n=11 and n=14 for the L1210 and FL5.12, respectively) gene ontology subsets (FDR<0.05, **Supplementary Figure 4, Supplementary Table 1, Methods**).

In order to determine the transcriptional profiles associated with various biophysical states in these cells, we ranked genes by how strongly their expression levels correlated with single-cell biophysical data (Spearman’s correlation coefficients, **Supplementary Table 1**). Both Spearman and Pearson correlation methods yielded similar results for all comparisons considered (**Supplementary Figure 3**). We then utilized the GSEA Preranked tool to determine which gene sets showed significant enrichment at either end of these ranked lists (FDR<0.05, **Methods, Supplementary Table 2**) [21]. For both cell types, genes ranked by correlation strength with single-cell mass were highly enriched for functional annotations relating to cell cycle progression (FDR<0.05, **Supplementary Table 2, Figure 2**). Specifically, genes related to early cell cycle events – such as DNA replication initiation – were more highly expressed in cells with lower masses whereas genes related to late cell cycle events – such as chromosome segregation – were more highly expressed in larger cells. Interestingly, both cell types revealed a larger number of genes that showed a significant positive correlation with mass relative to the number of genes with a significant negative correlation (**Supplementary Figure 4**).

Genes that showed significant correlation with cell mass in L1210 cells were significantly enriched amongst those previously shown to correlate with time since division, an alternative proxy for cell cycle progression, in the same cell line (FDR < 0.05, **Supplementary Figure 5, Supplementary Note 2**). Similarly, genes that showed a significant correlation between their expression levels and biophysical properties in FL5.12 cells were consistent when measured in a second, independent replicate of the linked biophysical and gene expression experiments (FDR<0.05, Supplementary Figure 5, Supplementary Note 2). These results demonstrate the quality and reproducibility of transcriptional measurements collected downstream of the sSMR.

To account for the linear relationship between mass and MAR in these cell types (ρ = 0.67 and ρ = 0.56 for L1210 and FL5.12, respectively, Figure 2), we focused our analysis on mass-normalized MAR, determined by dividing each cell’s MAR by its corresponding mass. This parameter describes a single cell’s growth efficiency, which is decoupled from mass-related confounders [18, 22]. For L1210 cells, genes ranked by strength of correlation between expression level and growth efficiency did not reveal any statistically significant enrichment of functional annotations (FDR>0.05). The FL5.12 cells, however, showed significant positive enrichment for functional annotations related to cell cycle progression amongst genes ranked by correlation strength with growth efficiency (FDR<0.05, **Supplementary Table 2**). Specifically, subsets of genes implicated in the G1-S transition and DNA replication showed a higher level of expression in cells of intermediate mass with the highest growth efficiencies (**Methods, Supplementary Figure 6**) [23]. These results are consistent with previous FL5.12 single-cell growth measurements, which revealed an increase in growth efficiency approaching the G1-S transition followed by a decrease later in the cell cycle [15].

### Characterizing CD8+ T cell activation with linked biophysical and gene expression measurements

Both the L1210 and FL5.12 models present stable distributions of biophysical and transcriptional profiles over the course of long-term propagation in bulk culture [24, 25]. However, one of the benefits of the sSMR platform is that it offers sufficient throughput to characterize cell populations that may be changing in their phenotypes over time. Primary CD8+ T lymphocytes are a prime example of this dynamic behavior, as they are known to drastically change their biophysical properties, transcriptional states, and metabolic characteristics in response to activation [17, 26, 27].

In order to characterize their response to activation, we collected single-cell biophysical and gene expression profiles for murine CD8+ T cells stimulated *in vitro* for either 24 or 48h (Figure 3**, Methods**). Although the cells for both time points displayed similar mass distributions, the cells measured after 48h of activation showed significantly higher growth efficiencies (P<0.001, Mann-Whitney U-test, Figure 3a,b). These two populations showed differential expression patterns consistent with T cell activation, including significant upregulation of Granzyme B (*Gzmb*) and IL-2 receptor (*Il2ra* and *Il2rb*) as well as significant downregulation of *Ccr7* in the 48h population compared to the 24h one (FDR<0.05, **Supplementary Table 3**). Similarly, gene set enrichment analysis performed on genes ranked by expression fold change between these time points revealed significant enrichment for gene sets related to immune cell effector function and glucose metabolism, consistent with functional and metabolic shifts that have been previously characterized in activated CD8+ T cells (FDR<0.05, **Supplementary Tables 4 and 5**) [26, 28].

**Figure 3.**
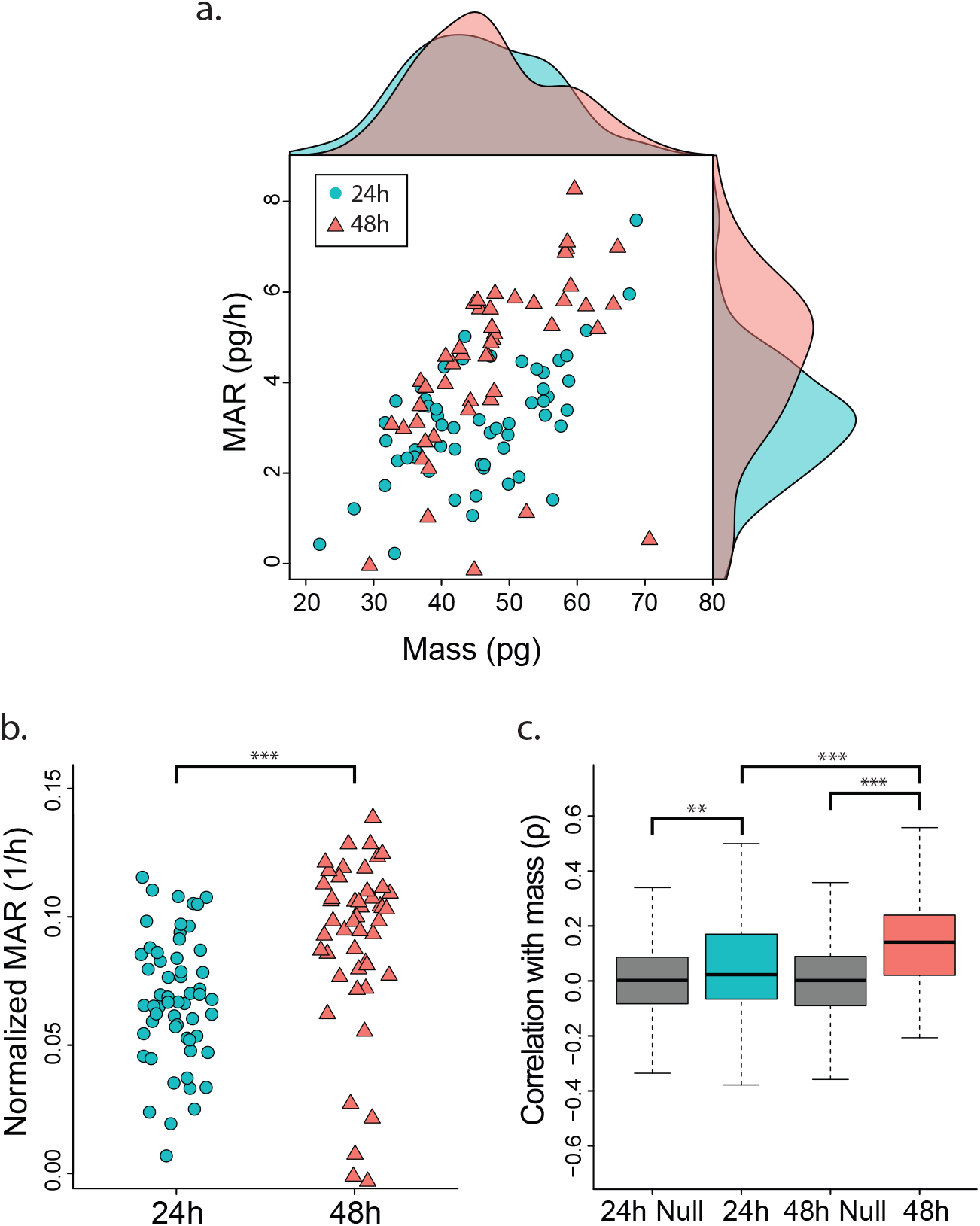
Linked biophysical and gene expression measurements of activated murine CD8+ T cells. (**a**) Plot of mass accumulation rate versus buoyant mass for murine CD8+ T cells after 24 h (blue points, n=59) or 48 h (red triangles, n=49) of activation *in vitro.* Kernel density plots, using the same color scheme, are included on the margins for both populations. (**b**) Plot of mass-normalized single-cell growth rates (growth efficiency) for the same murine CD8+ T cells activated for 24 or 48 hours *in vitro.* Groups were compared with a Mann-Whitney U-test (*** P < 0.001). (**c**) Box charts showing the Spearman correlation coefficients between single-cell mass measurements and the expression of a subset of genes previously found to be related to cell cycle in activated CD8+ T cells (300 genes) for cells activated for 24 or 48 hours. For comparison, the distribution of Spearman correlation coefficients for the same subset of cells after randomly assigning single-cell mass measurements is shown for each time point (**Methods**). Groups were compared with a Mann-Whitney U-test (*** P<0.001, ** P<0.01).

Cells activated for 48h also displayed a higher expression of genes related to protein production, including those involved in translation initiation and cytosolic ribosome activity (**Supplementary Table 5**). Araki et al. recently demonstrated a similar trend, noting an increase in translation activity over of the course of early T cell activation, as cells become more proliferative [29]. The measurements presented here suggest that this increase in translation activity is accompanied by, and potentially is tied to, an increased growth efficiency observed at 48h compared to 24h. This population-level relationship between growth efficiency and translation-related gene expression was also observable at the single-cell level for cells activated for 48h. Within this time point, genes ranked by correlation strength with single-cell growth efficiency once again showed significant enrichment for functional annotations relating to translation machinery (FDR<0.05, **Supplementary Table 2**). Despite a similar number of genes showing a significant correlation with growth efficiency in the 24h time point, these genes did not show any significant functional enrichment when ranked by correlation strength (FDR>0.05, **Supplementary Figure 4**). This result suggests that the coordination between single-cell growth efficiency and translation-related gene expression occurs later in the course of T cell activation.

The 48h time point revealed a greater number of genes that showed a significant correlation between expression level and cell mass relative to the 24h time point (**Supplementary Figure 4**). When determining the functional role of genes ranked by expression correlation with single-cell mass, only the 48h time point demonstrated significant cell cycle functional enrichment (FDR<0.05, **Supplementary Table 2**). Furthermore, a previously described set of genes known to correlate with an activated CD8+ T cell’s time since division – a proxy for cell cycle progression – showed a significantly stronger positive correlation with cell mass in the 48h population relative to 24h population (P<0.001, Mann-Whitney U-test, Figure 3) [25]. It is important to note that the 24 and 48h time points primarily capture cells before and after their first division event, respectively [30]. Although cells are accumulating mass, or “blasting”, in the first 24h, it is not until roughly 30 hours that cells undergo their first division and begin increasing in number and cycling in the traditional sense [30, 31]. Taken together, these results suggest that the level of coordination between cell cycle gene expression and cell mass increases later in T cell activation as cells begin actively dividing.

## Conclusion

The platform presented here enables linked measurements of single-cell biophysical properties and gene expression. We have demonstrated the feasibility of resolving distinct transcriptional signatures associated with subtle differences in single-cell mass and growth rate for stable L1210 and FL5.12 cell lines as well as for activated, proliferating CD8+ T cells. While the primary focus of this work was on conducting scRNA-seq downstream of the sSMR, we also envision this platform being a useful tool for linking biophysical data with other recently developed approaches that enable DNA sequencing, epigenomic characterization, or multi-omic measurements of single cells [6, 7, 32].

We believe that these linked measurements will offer a novel means of exploring a range of biological questions. For instance, when paired with recently developed computational approaches, these linked biophysical and transcriptional measurements may offer unprecedented insights into cell cycle regulation as well as provide an additional approach for addressing the potentially confounding effects of cell cycle in scRNA-seq analyses [33]. Clinically, mass and MAR have proven to be effective biomarkers for characterizing cancer cell drug susceptibility at the single-cell level [18, 22]. The ability to link these biophysical measurements with gene expression profiling offers the opportunity to move beyond the classification of responding and non-responding cells, and begin to explore the molecular mechanisms for such behaviors.

## Methods

### Cell culture and primary cell preparation

L1210 murine lymphocytic leukemia cells (ECACC) were cultured in RPMI 1640 (Gibco) with 10% fetal bovine serum and 1% antibiotic-antimycotic (Gibco). FL5.12 murine pre-B cells (Vander Heiden Lab, MIT) were cultured in the same media with the addition of 10 ng/ml IL-3 (R&D Systems). For all growth and collection experiments, cells were passaged to a concentration of 5 × 10^5^ cells/ml the night before to ensure consistent culture confluence at time of measurement. Naïve, CD8+ T cells were isolated from a 13 week old, male, C57BL/6J mouse. Splenocytes were subject to red blood cell lysis with ACK buffer (Gibco) followed by naïve CD8+ T cell isolation using a MACS-based isolation kit (Miltenyi Biotec). Purified cells were cultured in RPMI 1640 (Gibco) with 10% fetal bovine serum, 55 μM 2-mercaptoethanol (Gibco), 1% antibiotic-antimycotic (Gibco) and 100 U/ml IL2 (Peprotech). The naïve CD8+ T cells were activated *in vitro* with 5 μg/ml plate-bound anti-mouse CD3 (clone: 145-2c11, BioLegend), 0.5 μg/ml plate-bound ICAM-1/CD54 (R&D Systems), and 2 μg/ml soluble anti-mouse CD28 (clone: 37.51, BioLegend). Cells were seeded at a concentration of 1 × 10^6^ cells/ml in a 96 well plate and activated for either 24 or 48h prior to measurement in the sSMR.

Animals were cared for in accordance with federal, state and local guidelines following a protocol approved by the Department of Comparative Medicine at MIT (protocol number 0317-022-20).

### Single-cell growth measurements and collection

For all experiments, cells were adjusted to a final concentration of 2.5 × 10^5^ cells/ml to load single cells into the mass sensor array as described in **Supplementary Note 1**. Single-cell growth measurements were conducted as described previously [17]. In order to exchange buffer and flush individual cells from the system, the release side of the device was constantly flushed with PBS at a rate of 15 μL per minute (**Supplementary Figure 1**, P2 to P4). Upon detection of a single-cell at the final cantilever of the sSMR, as indicated by a supra-threshold shift in resonant frequency, a set of three-dimensional motorized stages (ThorLabs) was triggered to move a custom PCR-tube strip mount from a waste collection position to a sample collection position. The location of these motors was written to a file for the duration of the experiment in order to annotate single-cell mass and MAR measurements with well position, and thus transcriptional profiles, downstream. Each cell was collected in 5 μl of PBS directly in to a PCR tube containing 5 μl of 2X TCL lysis buffer (Qiagen) with 2% v/v 2-mercaptoethanol (Sigma) for a total final reaction volume of 10 μl. After each 8-tube PCR strip was filled with cells, the strip was spun down at 1000g for 30 seconds and placed immediately on dry ice. Following collection, samples were stored at −80 C prior to library preparation and sequencing.

### scRNA-Seq

Single-cell RNA isolation, cDNA library synthesis, next generation sequencing, read alignment and gene expression estimation were performed as described previously [34]. Briefly, Smart-Seq2 whole transcriptome amplification and library preparation were performed on single-cell lysates collected with the sSMR [35]. Single-cell libraries were then sequenced on a NextSeq500 using 30-bp paired end reads. Cells that exceeded a preliminary complexity threshold (4,000 genes for L1210 and FL5.12 cells, 2,000 genes for CD8+ T cells) and had successfully paired biophysical measurements were selected for further analysis. Overall, this yielded 85 out of 96 total L1210 cells, 124 out of 192 total FL5.12 cells, and 108 out of 192 total CD8+ T cells. These cells selected for analysis were sequenced to an average depth of 1,698,879 + 106,027 (s.e.m.), 760,919 + 36,679 (s.e.m.), and 1,333,686 + 90,744 (s.e.m.) reads for the L1210, FL5.12, and CD8+ T cells respectively. Reads were aligned using TopHat2 and expression estimates (transcripts per million; TPM) for all UCSC-annotated mouse genes (mm10) were calculated using RNA-seq by expectation maximization (RSEM) [36, 37]. The average transcriptome alignments were 67.4 + 0.38 % (s.e.m.), 64.8+ 0.51 % (s.e.m.), and 57.3 + 1.36 % (s.e.m.) for the L1210, FL5.12, and CD8+ T cells respectively. The average number of genes detected was 7,207 + 94 (s.e.m.), 6,891 + 81 (s.e.m.), and 5,149 + 159 (s.e.m.) for the L1210, FL5.12, and CD8+ T cells respectively (**Supplementary Figure 2**).

### Gene expression analysis

All analysis was performed on log-transformed expression level measurements (ln(TPM+1)). Data preprocessing was conducted with the Seurat package for R [10]. All genes that were detected in >5% of cells were included in the final analysis for each group of cells (L1210, FL5.12, and CD8+ T cells).

Ranked gene lists were created for each cell population by determining the gene-wise correlation coefficient (Spearman) between log-transformed gene expression level and either single-cell mass or growth efficiency (**Supplementary Table 1**). Spearman and Pearson correlation coefficients yielded similar results for all conditions measured (**Supplementary Figure 3**). Gene set enrichment was computed for these ranked lists using the GSEA Preranked tool, implemented with the fgsea package in R (**Supplementary Table 2**) [21, 38].

Differential expression analysis for the 24 versus 48h CD8+ T cell measurements was performed using the FindMarkers function of Seurat with the Wilcoxon rank sum test (**Supplementary Table 3**). Genes were also ranked by log-normalized fold-change expression difference between the 24 and 48h time points and analyzed with the GSEA Preranked tool (**Supplementary Tables 4 and 5**).

To define the null distribution of correlation coefficients described in Figure 3, we determined the Spearman correlation between cell cycle gene expression levels and mass for randomly shuffled data sampled from the experimental values (i.e. mismatched single-cell mass and gene expression data). After 10,000 iterations of this process, we found the average mean and standard deviation values of these correlation coefficient distributions which were used to define the null distributions presented.

## Abbreviations

scRNA-seq: : single-cell RNA-sequencing;
sSMR: : serial suspended microchannel resonator;
MAR: : mass accumulation rate;
FDR: : false discovery rate

## Author contributions

R.J.K., N.L.C., S.O., N.C., and S.R.M. designed and implemented the platform. R.J.K., A.K.S., and S.R.M. designed the experiments. R.J.K., A.J.G., and N.L.C. performed the sSMR experiments. S.M.P., A.J.G., and R.D. performed scRNA-seq. R.J.K., S.M.P., A.J.G., and M.M.S. analyzed the data. R.J.K., A.K.S., and S.R.M. wrote the manuscript, with input from all authors.

## Acknowledgements

This work was supported by Cancer Systems Biology Consortium U54 CA217377 (SRM and AKS), R33 CA191143 (SRM), the Searle Scholars Program (A.K.S.), the Beckman Young Investigator Program (A.K.S.), NIH New Innovator Award 1DP2OD020839 (A.K.S.), NIH 5U24AI118672 (A.K.S.), NIH 1R33CA202820 (A.K.S.), NIH 2U19AI089992 (A.K.S.), NIH 1R01HL134539 (A.K.S.), NIH 2RM1HG006193 (A.K.S.), 2P01AI039671 (A.K.S.), and partially by Cancer Center Support (core) Grant P30-CA14051 from the National Cancer Institute.

## Data access

Raw and processed scRNA-seq data will be uploaded to the Gene Expression Omnibus prior to publication. All SMR data, hardware interface code, and data analysis code are available upon reasonable request.

## Conflict of interest statements

R.J.K., M.M.S., S.O., and S.R.M. declare competing financial interests as cofounders of Travera, which develops technology relevant to the research presented. S.R.M. declares competing financial interests as a cofounder of Affinity Biosensors, which develops technology relevant to the research presented.

## Supplementary Figure Captions

**Supplementary Figure 1 |** Fluidic regimes for maintaining cell spacing in the sSMR

(**a**) Schematic of sSMR presented in Figure 1 denoting the array entrance region used for fluidic simulation presented in (**b**) (dashed box outline). (**b**) COMSOL fluidic simulations demonstrating the loading (left) and flushing (right) fluidic regimes described in Supplementary Note 1. The imaging region used to trigger between each fluidic state is outlined (solid box).

**Supplementary Figure 2 |** Quality metrics for scRNA-seq libraries

Violin plots and overlaid points showing the number of genes detected (left), sequencing depth (center), and transcriptome alignment (right) for each scRNA-seq library prepared for (**a**) L1210 cells, (**b**) FL5.12 cells and (**c**) CD8+ T cells activated for either 24 or 48h (blue and red outlines, respectively) that passed initial quality thresholds and were used for further analysis (**Methods**).

**Supplementary Figure 3 |** Comparison of Pearson and Spearman coefficients for correlations between gene expression and biophysical parameters

Plots of the Pearson coefficient versus Spearman coefficient for expression level correlations with either mass (left column) or mass-normalized MAR (right column) for L1210, FL5.12, and CD8+ T cells (24 and 4h time points). Each cell type lists the total number of genes being compared and each plot indicates the Spearman coefficient between the Spearman and Pearson coefficients across all genes. Each measurement set reveals similar gene-level rankings for both Spearman and Pearson coefficients.

**Supplementary Figure 4 |** Expression level correlation with biophysical parameters for L1210, FL5.12, and CD8+ T cells

Bar plots denoting the correlation strength of individual gene’s expression levels with either mass (left) or mass-normalized MAR (right) for (a) L1210 cells (n = 11,469 genes), (b) FL5.12 cells (n = 11,040 genes), (c) CD8+ T cells after 24h of activation (n = 9,015 genes), and (d) CD8+ T cells after 48h of activations (n = 9,015 genes). Genes are plotted in rank order where genes with highest positive and negative correlations with biophysical parameters are found at the left-most and right-most portion of the x axis, respectively. For each data set, a null distribution of correlation coefficients was determined by finding the correlation between gene expression and mass for randomly permuted data. After 10 iterations, we determined the average standard deviation of these distributions of correlation coefficients. Any individual gene that had a correlation coefficient with an absolute value greater than twice the standard deviation (P<0.05, denoted by the dashed lines in the plots) was considered significant (red bars), all genes presented as blue bars fell below this threshold. The number of genes showing a significant positive or negative correlation with the biophysical parameter of interest are shown in each plot.

**Supplementary Figure 5 |** Reproducibility of linked measurements for L1210 and FL5.12 cells (**a**) Enrichment plots for genes with significant positive (left, n = 1,166) or negative (right, n = 134) correlations with single-cell mass amongst genes ranked by expression level correlation with time since division in L1210 cells – determined by Kimmerling et al. (**Supplementary Note 2**) [25]. Significant enrichment (FDR = 0.0009 and 0.0003 for positive and negative sets, respectively) suggests that a consistent cell cycle gene expression signature correlates with both cell mass and time since division in L1210 cells. (**b**) Enrichment plots for genes with significant positive (left, n = 874) and negative (right, n = 191) correlations with FL5.12 cell mass amongst a full gene list ranked by expression level correlation with FL5.12 cell mass from a second, independent experiment. The significant enrichment here (FDR = 0.0004 and 0.0016 for positive and negative sets, respectively) demonstrates a reproducible gene expression signature corresponding to FL5.12 mass. (c) Same analysis as in (b) for genes that correlated significantly with mass-normalized growth rate (growth efficiency, n = 309 and 621 genes for positive and negative correlations, respectively) as opposed to mass, demonstrating reproducible growth-related gene expression signatures as well (FDR = 0.0002 and 0.0205 for positive and negative sets, respectively).

**Supplementary Figure 6 |** Cell cycle gene expression versus growth rate in FL5.12 cells

Plot of mass versus mass-normalized growth rate (growth efficiency) for a subset of the FL5.12 cells depicted in Figure 2 that were captured downstream for scRNA-seq (n = 124). Points are colored by G1/S score rank with highest expression score corresponding to red and lowest expression score corresponding to blue. The “cell cycle G1/S phase transition” gene ontology term was found to be significantly enriched amongst genes ranked by correlation with growth efficiency. To determine the G1/S transition scores for single FL5.12 cells we found the average of mean-centered, z-score scaled expression values for the leading-edge genes of the “cell cycle G1/S phase transition” gene ontology term – these are the genes that, when included, give rise to the highest enrichment score for this term (n = 40 genes) [21].

## Supplementary Notes

**Supplementary Note 1 |** Maintaining minimum cell spacing in mass sensor array

Loading single cells into the mass sensor array at a fixed, minimum spacing requires the implementation of active switching between two distinct fluidic states. Initially, equivalent pressures are applied to the upstream and downstream ports on the bypass channel leading in to the array (**Supplementary Figure 1**, ports P1 and P3). In this “loading” configuration, all streamlines are directed into the array and therefore cells in the bypass channel will enter the array. An imaging region at the entrance to the mass sensor array (outlined in **Supplementary Figure 1**) is used as an indication of when a cell has been successfully loaded. Real-time optical peak detection within this region is used to switch from this loading fluidic state to a “flushing” regime wherein the upstream pressures (P1) is increased and the downstream pressure (P3) is decreased such that a vast majority of streamlines continue along the bypass channel with a small fraction entering the array. Because cells are of finite size and occupy several streamlines, they are directed along the bypass channel and not drawn in to the array. Importantly, during this process the pressure at the entrance to the mass sensor array is maintained at a fixed value, therefore any cells that have entered the array continue to flow at a constant speed. Therefore, although the volumetric flow rate is maintained across the array while flushing, no additional cells are loaded. After a desired amount of time has elapsed the system is automatically returned to the loading configuration to obtain the next cell for measurement.

**Supplementary Note 2 |** Determining reproducibility of gene signatures related to mass and MAR

In order to determine the reliability and reproducibility of the linked biophysical and gene expression profiles, it was important to compare these signatures with additional results collected from independent experiments. For L1210 cells, single-cell gene expression profiles had previously been collected for cells with known times since division (TSD), a proxy for cell cycle progression [25]. We therefore hypothesized that the list of genes with expression levels that correlated significantly with single-cell mass (an alternative proxy for cell cycle progression) would show significant overlap with genes that correlated strongly with TSD. To determine the extent of this similarity, we constructed two test gene sets for gene set enrichment analyses: one which included genes with a significant positive correlation with cell mass and another which included genes with a significant negative correlation with cell mass (**Supplementary Figure 5a, Supplementary Table 2**). These gene subsets were compared to the full L1210 gene list measured previously, with genes ranked by how strongly their expression levels correlated with TSD. Genes with a significant positive correlation with mass were significantly over-represented amongst genes that showed a positive correlation with TSD in prior measurements (FDR<0.05). Similarly, genes with a significant negative correlation with mass were significantly over-represented amongst genes that showed a negative correlation with TSD (FDR<0.05). These results indicate that similar sets of genes are correlated with both TSD and single-cell mass, suggesting consistency between the measurements collected here and those collected previously.

Next, we sought to perform a similar comparison for FL5.12 cells. However, in contrast to L1210 cells, no single-cell gene expression measurements had been collected for these cells previously. We therefore conducted a second, independent experiment where single-cell mass and MAR measurements were collected upstream of scRNA-seq for FL5.12 cells (**Supplementary Figure 5b,c**). Using this independent data set, we generated full gene lists that were ranked by correlation strength with either mass or mass-normalized MAR. Then we once again constructed test gene sets, this time containing genes from the original FL5.12 data set with significant correlations (both positive and negative) with either mass or mass-normalized MAR (P<0.05). Following the same analysis described above, we found that gene sets correlating with both mass and mass-normalized MAR showed significant overlap between both replicate experiments (FDR<0.05). This once again demonstrates the reproducibility of the gene expression signatures that correlate with single-cell biophysical properties.

## Supplementary Material

**Supplementary Table 1 |** Gene lists ranked by correlation with either mass or mass-normalized MAR for L1210, FL5.12, and CD8+ T cells (24 and 48h activations) with corresponding Spearman correlation coefficients. Genes that are either significantly positively or negatively correlated with the biophysical measurement of interest are highlighted in red.

**Supplementary Table 2 |** Gene set enrichment reports for all the ranked gene lists presented in **Supplementary Table 1**. Enrichments were generated using the fgsea tool in R. Only gene sets with a false discovery rate (FDR) value less than 0.05 are included.

**Supplementary Table 3 |** List of significantly differentially expressed genes between the 24 and 48h time points for the activated CD8+ T cells with corresponding FDR values and log-normalized fold change values. Negative values indicate genes expressed at a higher level in the 48h time point.

**Supplementary Table 4 |** CD8+ T cell gene list ranked by log-normalized fold change in gene expression between the 24 and 48h activation time points. Negative values indicate genes expressed at a higher level in the 48h time point.

**Supplementary Table 5 |** Gene set enrichment report for the ranked gene list presented in Supplementary Table 4. Enrichments were generated using the fgsea tool in R. Only gene sets with a false discovery rate (FDR) value less than 0.05 are included.

